# lme4breeding: a fast linear mixed model for multi-trait/multi-environment experiments

**DOI:** 10.1101/2024.05.17.594795

**Authors:** Giovanny Covarrubias-Pazaran

## Abstract

Mixed models are a cornerstone in quantitative genetics to study the genetics of complex traits. A standard quantitative genetic model assumes that the effects of some random factors (e.g., individuals) are correlated based on their identity by descent and state. In addition, other relationships arise in the genotype by environment interactions (i.e., covariance structures). Open-source mixed model routines are available but do not account for complex covariance structures or are extremely slow to fit big/dense genomic models. The lme4breeding R package was developed as an extension of the lme4 package and allows correlated random effects and complex covariance structures to be fitted for responses with different distributions. The correlation between levels of the random effect (e.g., individuals) is accounted for by eigen-decomposing the relationship matrix and post-multiplying the incidence matrix of the levels of this random factor by the Cholesky decomposition of the corresponding (co)variance matrix (eigen values) and the response by the eigen vectors. For big genomic models the eigen decomposition of relationship matrices coupled with sparse matrix solvers presents a massive increase in speed compared to dense formulations used in other popular software and the same level of accuracy is achieved.

## 1 Introduction

Mixed models are the cornerstone to estimate genetic parameters and predict surrogates of merit (i.e., breeding values, general combining ability, or total genetic value) associated with normally distributed traits (Henderson et al., 1959; Henderson, 1963, 1973). In linear mixed models with Gaussian distribution, the marginal likelihood has a closed form, and maximum likelihood or REML estimation can be performed, whereas in nonlinear models the likelihood does not have a closed form and approximations are needed. Generalized mixed models allow to fit different distributions from the exponential family of distributions (Tempelman, 1988).

The development of mixed model packages in the R environment (Team, 2008) has increased in the last decade due to a quick learning curve and easiness to code complex algorithms and sophisticated graphics. R is an open-source system written and maintained by volunteers (www.r-project.org). The **lme4** package (Bates, 2015) is the gold-standard when it comes to fitting linear models and GLMM to data. The program handles any number of random effects, nested or cross-classified, and uses a combination of sparse and dense matrix representations to process large data sets at high speed. The use of **lme4** for genetic analysis was limited because it does not allow covariances between levels of a random effect. To close that gap Vazquez et al. (Vazquez et al., 2010) developed a package called **pedigreemm** to expand lme4 capabilities to allow correlations between levels of random effects, specifically pedigrees and made it available on CRAN. With the advent of genomic information, Ziyatdinov et al. (Ziyatdinov et al., 2018) expanded the **lme4** to include genomic relationship matrices inspired by pedigreemm with a package called lme4qtl but unfortunately was neither published on CRAN nor properly documented. Camaal et al. (Caamal-Pat et al., 2021) published a package to do the exact same as lme4qtl and called it **lme4GS** which was recently published on CRAN but did not add anything new on top of what lme4qtl had developed. Any **lme4** extension has shown to be extremely slow to fit genomic relationship matrices due to the dense formulation when a relationship matrix exists.

In the present note, we discuss the approach implemented by **lme4breeding** for GLMM to include covariances between levels of random effects (any Cholesky factor), coupled with the eigen (**HDH**^*′*^) decomposition of the relationship matrix to speed up operations in balanced settings (e.g., multi-trait imputed data). In addition, customized random effects (e.g., overlay models for half-diallel or indirect genetic effects) can also be provided, and the ability to model complex covariance structures like the ones found in the genotype by environment problem (e.g., factor analytic structures). We show the use of the package with two examples using data included in the package. The package is available on CRAN (https://CRAN.R-project.org/package=lme4breeding).

## 2 Materials and methods

The classical generalized linear mixed model follows the formulation:

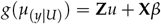

where y is the random variable representing the response; g(.) is a function that links the response with a model that is linear in the explanatory variables; *µ*_*y* |*U*_ = *E*[*y*| *U* = *u*] is the expectation of the response conditional to the random effects; *β* is the vector of fixed effects; **X** is the incidence matrix relating fixed effects to *gµ*_(*y*|*U*)_); u is the vector of random effects, which may row-bind multiple factors, where each factor of random effect is distributed as 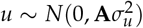, where N(.) represents the normal distribution with mean and variance indicated in the parentheses; 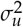 is a variance-covariance matrix; **A** is a relationship matrix; and **Z** is the incidence matrix relating u to *g*(*µ*_(*y* |*U*)_). In quantitative genetics, the variance-covariance matrix of individuals is modeled through additive and non-additive relationships based on the identical-by-descent and identical-by-state theory (Mrode, 2005).

To obtained variance components of this model, restricted maximum likelihood is commonly used (Patterson and Thompson, 1971). On the other hand, on the generalized linear mixed model, parameters from the exponential family of distributions are obtained by minimizing the Laplacian approximation to the deviance function as suggested by to Bates (Bates, 2015). A sparse implementation to solve these equations has been implemented in a popular package called **lme4**, which is the gold-standard in the R language.

Vazquez et al. (Vazquez et al., 2010) proposed that the methodology described by Harville and Callanan (Harville and Callanan, 1989) consisting of post-multiplying the model matrix **Z** by the Cholesky decomposition of the numerator relationship matrix (**A**) can be used to leverage the **lme4** machinery, ensuring that **A** is a positive-definite matrix (or positive semi-definite). Remember that **A** = **LL**^*′*^, where **L** is the Cholesky factor or decomposition. Subsequently, let **Z**^∗^ = **ZL**, then

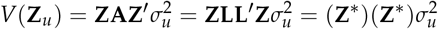

Define *u*^∗^ = **L**^−1^ *u*, and rewrite *g*(*µ*_(*y*|*U*)_) as:

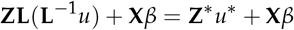

Assuming 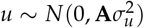, the distribution of u* is 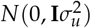 because

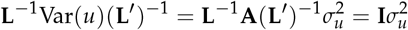

Making the elements of u* mutually independent. The **lme4** machinery can then be applied to *g*(*µ*_(*y*|*U*)_) = **Z**^∗^*u*^∗^ + **X***β* because the random effects are now independent.

In addition to that, to speed up the genomic models we applied the **HDH**^*′*^ decomposition proposed by Lee and Van der Werf (Lee and der Werf, 2016). The parameterization is:

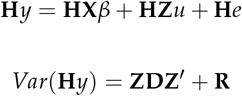

Where **H** and **D** are the eigen vectors and eigen values respectively of any relationship matrix (e.g., genomic relationship matrix). **H** can be derived as:

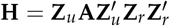

Where **Z**_*u*_ is the incidence matrix for the individuals, **Z**_*r*_ is the incidence matrix for the records for the individuals, and * is the element-wise multiplication of two objects (Schur product). For more technical details the reader is suggested to check the canonical publications from Harville and Callanan (Harville and Callanan, 1989) and Lee and Van der Werf (Lee and der Werf, 2016).

### Data examples

#### Dataset 1

Information from a collection of 599 historical CIMMYT wheat lines was used. The wheat data set is from CIMMYT’s Global Wheat Program. Historically, this program has conducted numerous international trials across a wide variety of wheat-producing environments. The environments represented in these trials were grouped into four basic target sets of environments comprising four main agroclimatic regions previously defined and widely used by CIMMYT’s Global Wheat Breeding Program. The phenotypic trait considered here was the average grain yield (GY) of the 599 wheat lines evaluated in each of these four mega-environments. Lines were genotyped using 1447 Diversity Array Technology (DArT) generated by Triticarte Pty. Ltd. (Canberra, Australia; http://www.triticarte.com.au). The DArT markers may take on two values, denoted by their presence or absence. Markers with a minor allele frequency lower than 0.05 were removed, and missing genotypes were imputed with samples from the marginal distribution of marker genotypes. The dataset and script can be accessed by typing `?DT_wheat` in the terminal.

#### Dataset 2

Information from a collection of 413 rice hybrids was used to show the advantages of a multi-trait fit. The data set belongs to the Rice Diversity Organization Program. The lines are genotyped with 36,901 SNP markers and phenotyped for more than 30 traits (Zhao et al., 2011). The dataset and script can be accessed by typing `?DT_rice` in the terminal.

Both datasets represent a balanced scenario where the eigen (**HDH**^*′*^) decomposition of the relationship matrix can be used to convert the problem into a sparse problem where the **lme4** machinery has its strength. Also unbalanced datasets can be fitted but the eigen decomposition ca not be used, slowing down the computations.

### Fitting the models

#### Dataset 1

The wheat yield lines (dataset 1) were analyzed with the classical multi-environment GBLUP model:

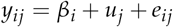

Where *y*_*ij*_ is the observation (yield performance) of individual j in location i; *β*_*i*_ is the fixed effect associated to location i (i=1,…,4), *u*_*j*_ is the breeding value of individual j (j=1, …, 599), and *e*_*ij*_ is the random residual associated to the observation. The following distributions are assumed for the vector of breeding values and residual effects.

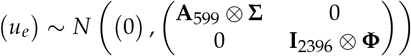

Where u is the vector of breeding values and e is the vector of residual effects. The **A** matrix represents the matrix of additive relationships between wheat lines, whereas ⊗ denotes the Kronecker product of two matrices. The variance-covariance of breeding values Σ is:

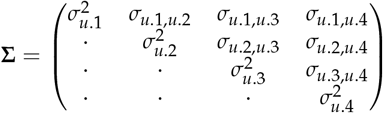

Representing the genetic covariance between the four locations and Φ represents a one-by-one matrix of residual variance. This model was fitted in R as follows:

~~~
system.time(
 mix1 <- lmebreed(pheno ∼ (0 + e1 + e2 + e3 + e4 | id),
           relmat = list(id=K),
           rotation = TRUE,
           verbose = 0L, trace = 0L,
           data=DT)
)
#> user system elapsed
#> 0.316 0.004 0.321
~~~

The mix object contains the fitted model and is of class lmebreed. The left-hand side of the formula denotes the response variable and the right-hand side the explanatory variable. Terms in parentheses represent the random effects [e.g. (intercept | slope)] expressed as random regressions (slopes) over numerical covariates (intercepts) as normally done in **lme4**. In this model the columns e1 to e4 are columns coded with a 1 if the record belongs to the ith environment and a 0 when it does not. These random intercepts are written on the left side of the | symbol. The term id represents the random slope being regressed over the environment covariates (e1 to e4), in this case the wheat line evaluated.

The parameter relmat is a named list containing the relationship matrices. The symbol “|” represents a unstructured covariance between the random slopes. The parameter rotation controls the activation of the eigen decomposition. The function `system.time()` is used to measure how long the procedure takes to run the analysis. As the result shows, it took less than a second for this example, run on a Mac-OS system with an M1 chip and 8 GB of RAM memory.

#### Dataset 2

This example uses a multi-trait linear model. The rice dataset has been modeled previously with univariate models for genome-wide association studies (Zhao et al., 2011). The multi-trait GBLUP model can be defined as:

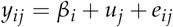

Where *y*_*ij*_ is the observation of the jth individual for the ith trait value; *β*_*i*_ is the fixed effect of a trait intercept (i=1,…,10), *u*_*j*_ is the breeding value of hybrid j (j=1, …, 413), and *e*_*ij*_ is the random residual associated to the observation. Trait observations are arranged in the long format to suit the **lme4breeding** machinery by creating a column for each of the 10 traits fitted. The following distributions are assumed for the vector of breeding values and residual effects.

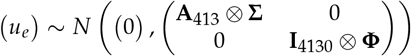

Where u is the vector of breeding values and e is the vector of residual effects. A represents the matrix of additive relationships between the rice hybrids, whereas ⊗ denotes the Kronecker product of two matrices. The variance-covariance Σ for the multi-trait fit is:

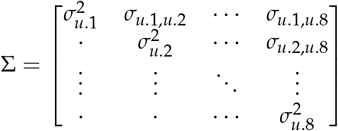

Representing the genetic covariance between the eight traits and Φ represents a one-by-one matrix of residual variance. This model was fitted in R as follows:

~~~
system.time(
 mix2 <- lmebreed(value ∼ trait +
                             (0+Flowering.time.at.Arkansas + Culm.habit +
                             Flag.leaf.length + Panicle.number.per.plant +
                             Plant.height + Panicle.length + Seed.number.per.panicle +
                             Florets.per.panicle + Seed.length + Seed.width | geno),
                  relmat = list(geno=A),
                  control = lmerControl(
                   optCtrl = list(maxfun = 2000, maxeval = 2000)
                   ),
                  rotation = TRUE, verbose = 0L, trace = 0L,
                  data=DTL)
)
#> user system elapsed
#> 8.501 0.094 8.619
~~~

In this model the column `trait` is coded as factor so **lme4breeding** creates internally ten indicator columns (random intercepts), one for each trait that associate observations to traits. The random regression of genotypes proceeds over traits with an unstructured covariance (| symbol). The term geno represents the random factor (slope) being regressed over the traits, in this case the rice hybrid evaluated. The parameter `relmat` is the additive relationship matrix between hybrids. The parameter `rotation` controls the activation of the eigen decomposition. The function `system.time()` was used to measure how long the procedure took. As the result shows, it took about 10 seconds for this example, run on a Mac-OS system with an M1 chip and 8 GB of RAM memory for 1000 iterations of the non-linear optimizer (nloptwrap).

## 3 Results

This section will show how to extract results from lmebreed objects.

**Figure 1:**
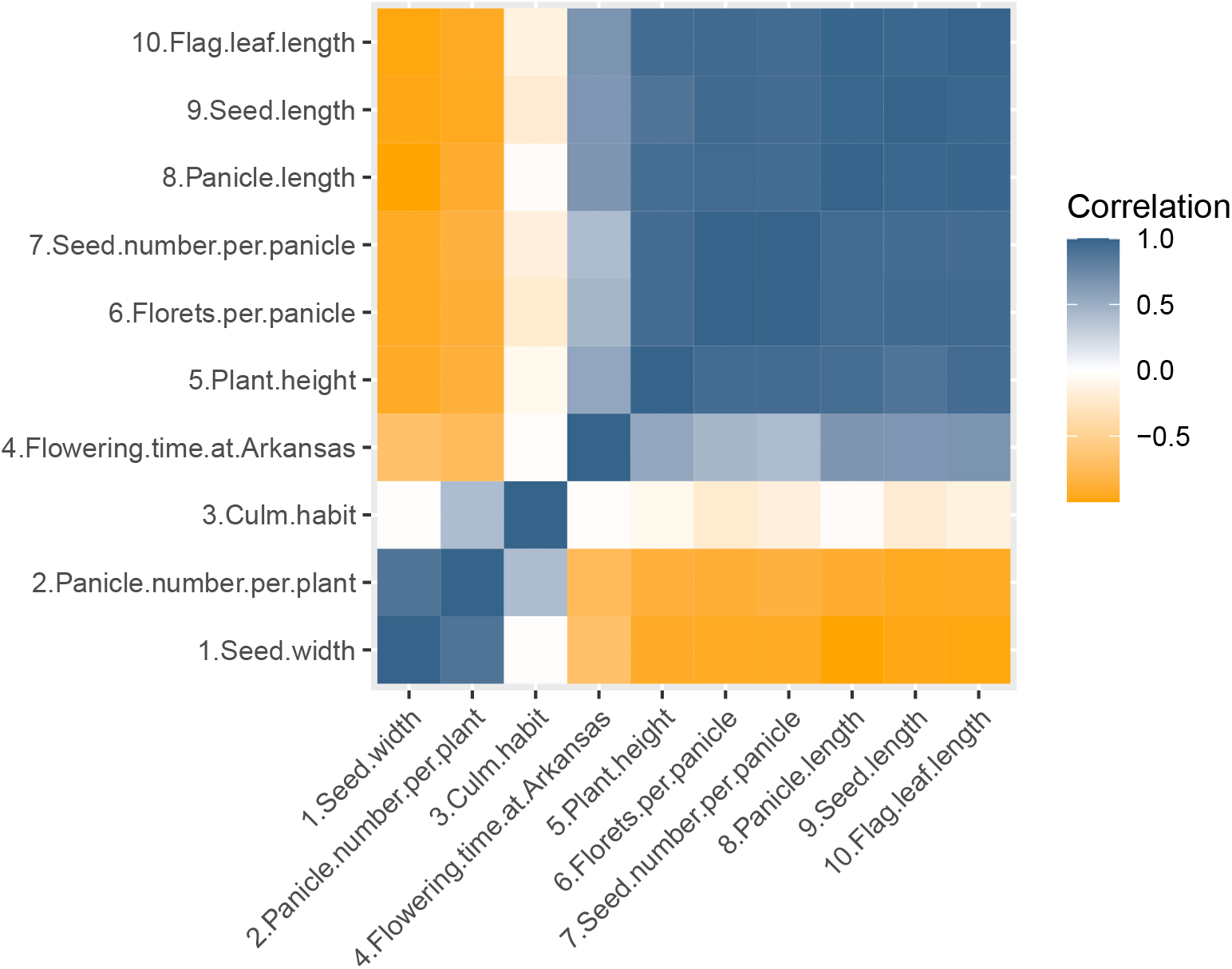
Visualization of the REML estimates of the genetic variance covariance matrix transformed into correlations from the dataset number 2 where an unstructured model for 10 traits was fitted.

### Variance components

The variance components associated to the random effects can be observed calling the `summary()` function but they can be retrieved more easily using the `varCorr()` function as follows:

~~~
vc1 <- VarCorr(mix1); print(vc1)
#> Groups  Name Std.Dev. Corr
#> id       e1  0.58334
#>          e2  0.73484   -0.180
#>          e3  0.70790   -0.275 0.994
#>          e4  0.62182   -0.349 0.759 0.808
#> Residual     0.69739
~~~

This function already returns the variance covariance matrices for each random effect in a ready-to-use format where each element of the list is the variance-covariance matrix for each random effect. The one above is the unstructured variance covariance matrix of the multi-environment model. The residual variance can be retrieved in the same way than **lme4** since it is stored in the `varCorr()` object as the sc attribute:

~~~
attr(vc1, “sc”)^2
#> [1] 0.4863531
~~~

### Random and fixed effects

Effects for the fixed and random factors are of special interest for geneticists to estimate merit of individuals, allelic effects, and test other evolutionary hypotheses. These effects can be retrieved from the lmebreed objects using the `fixef()` and `ranef()` functions. The fixed effects can be retrieved for the dataset 1 as:

~~~
fixef(mix1)
#> (Intercept)
#> -2.418247e-16
~~~

The random effects retrieved by the `ranef()` function are returned in their original scale (u). To calculate u, the transformed predicted effects inside the model (*u*^∗^ = **L**^−1^)*u*) are premultiplied/transformed by **L**, the Cholesky factor of the relationship matrix which are stored in the lmebreed object (*u* = **L***u*^∗^). If the eigen decomposition (**HDH**^*′*^) is activated using the rotation argument, an additional rotation/transformation is performed (*u*^∗∗^ = **H**^*′*^(**L**^−1^)*u*) to recover the real effects by premultiplying **L***u* by **H** (*u* = **HL***u*^∗∗^), the eigen vectors of the relationship matrix. The function is used as follows:

~~~
head(ranef(mix1)$id)
#>                e1         e2        e3          e4
#> 103122  -0.6929415 -2.1007523 -1.9158147 -1.3705767
#> 1047493  0.1631503  0.2057433  0.2054566  0.4112676
#> 1058137 -0.5149879 -2.0334580 -1.8639664 -1.2678927
#> 1064007  0.4152701  0.7155706  0.6379097  0.4966868
#> 1066175  0.4349651 -0.2379518 -0.2713296 -0.1952754
#> 1066279  0.2008866  1.2076310  1.1088687  0.6672886
~~~

In the table above rows are the levels of the random effect and columns are the estimate at each intercept level in the random regression (e.g., environment in the wheat example and traits in the rice example).

The standard errors of the random effects are of special interest for scientist to carry on comparisons and hypothesis testing. Just like with random effects, the recovery of accurate standard errors requires to pre and post multiply the inverse of the coefficient matrix by the eigen vectors when the rotation is used but they do not require to be multiplied by the Cholesky factor **L**. To retrieve the standard errors the argument condVar in the `ranef()` function can be set to `TRUE` as follows in the example dataset 1:

~~~
attr(ranef(mix1, condVar=TRUE)$id, which="postVar")[,,1]
#>           [,1]      [,2]      [,3]      [,4]
#> [1,] 0.7972351 0.5636428 0.7383110 1.0410957
#> [2,] 0.5207929 0.9235207 0.4821591 0.9323149
#> [3,] 1.0856695 0.4817336 0.8027438 1.0367386
#> [4,] 0.4690776 0.7445102 0.7943130 1.0905356
~~~

Where the attribute `postVar` retrieves the standard errors for each level of the random effect and each slice corresponds to one level for the covariance structure specified (unstructured between four environments).

### Time comparisons against other popular software

In order to create an intuition of the time gains using **lme4breeding** compared to popular software, we run the example dataset 1 that includes 599 lines genotyped and tested in 4 environments modeled with an unstructured covariance matrix using the following libraries; `lme4qtl (0.2.2), sommer (4.3.4)`, and `Asreml (4.2)` using the default parameters on a Mac-OS machine with an M1 chip and 8 GB of RAM memory. The results were the following:

As it can be seen in Table 1 the ten variance-covariance components were almost identical among the libraries except for `sommer` that had problems to converge for this specific dataset. This shows that the accuracy of **lme4breeding** to estimate variance components is comparable to other popular software since it is based on the **lme4** package which is considered a robust mixed model machinery.

**Table 1:** Variance-covariance components obtained by different libraries for the example dataset 1 that includes 599 genotypes tested in 4 environments of data fitted as an unstructured model with a genomic relationship matrix.

| Package | $\sigma^2_{e1}$ | $\sigma^2_{e2}$ | $\sigma^2_{e3}$ | $\sigma^2_{e4}$ |
| --- | --- | --- | --- | --- |
| lme4breeding with rotation | 0.34 | 0.53 | 0.50 | 0.38 |
| lme4breeding w/o rotation | 0.34 | 0.54 | 0.50 | 0.38 |
| lme4qtl | 0.34 | 0.54 | 0.50 | 0.38 |
| sommer | 0.35 | 0.16 | 0.15 | 0.12 |
| Asreml | 0.34 | 0.54 | 0.51 | 0.38 |

**Table 2:** Time comparison in seconds (average across 10 samples) between different libraries for a genotype (599) by environment (4) problem fitted with an unstructured variance-covariance structure.

| Package | User_time | System_time | Elapsed_time | X_times |
| --- | --- | --- | --- | --- |
| lme4breeding with rotation | 0.702 | 0.048 | 0.772 | 1 |
| lme4breeding w/o rotation | 3813.413 | 25.963 | 3838.229 | 5432 |
| lme4qtl | 3666.069 | 0.567 | 3665.001 | 5222 |
| sommer | 451.330 | 5.296 | 454.727 | 643 |
| Asreml | 470.000 | 79.040 | 171.940 | 670 |

**Table 3:** Time comparison in seconds (average across 10 samples) between different libraries for a genotype (413) by trait (10) problem fitted with an unstructured variance-covariance structure in the rice dataset and a genomic relationship matrix.

| Package | User_time | System_time | Elapsed_time | X_times |
| --- | --- | --- | --- | --- |
| lme4breeding with rotation | 3.321 | 0.102 | 3.483 | 1 |
| Asreml | 988.830 | 152.740 | 526.470 | 298 |

In addition, the comparison of times with the different software shows the great advantage of the eigen decomposition trick over the dense formulation over both, **lme4breeding** without rotation and other popular software, ranging from ∼300x to >5000x for an unstructured model in the two datasets shown in this manuscript. The gain is even higher when the problem becomes denser. This increase in speed is thanks to the coupling of the eigen decomposition proposed by Lee and Van der Werf (Lee and der Werf, 2016) with the Cholesky decomposition of Harville and Callanan (Harville and Callanan, 1989) to create a totally sparse formulation of the genetic evaluation problem and using the **lme4** machinery that has been optimized for sparse problems (Bates, 2015). These two methods together achieve great speeds for genetic evaluation difficult to match with the dense formulation software. The major drawback of the eigen-decomposition of the relationship matrix is that only one relationship matrix can be eigen-decomposed, and the rest of the random effects can only have a diagonal relationship matrix (Lee and der Werf, 2016). In addition, this only applies for balanced datasets as pointed out by Lee and Van der Werf (Lee and der Werf, 2016). Fortunately, imputing the missing records in unbalanced datasets with simple methods (e.g., mean or median) gives highly correlated BLUP estimates (>0.98) compared to the dense formulation. In addition, many of the genetic evaluation problems only have one dense relationship matrix fitted and in multi-trait and multi-environment problems (the focused of this research) we can use the imputation trick to be able to use the eigen decomposition. Other models like factor analytic, diagonal, compound symmetry, and independent residual models are also possible to fit in **lme4breeding**, and the documentation included in the package provides many additional examples to know how to fit these and others.

## 4 Conclusions

The **lme4breeding** package enables the use of the eigen decomposition of the relationship matrix coupled with the Cholesky decomposition, resulting in a diagonal (co)variance structure of the random effects that speeds up the REML computation, especially in multi-trait or multi-environment scenarios that are balanced. This approach expands the powerful **lme4** capabilities to fit random regressions and sparse problems to solve many of the plant and animal breeding problems of genetic evaluation. As shown, hundreds of genotyped individuals tested in many environments or many traits can be fitted in few seconds due to the sparse parameterization of the problem and surpasses the capabilities of similar software. The tool is available in R (https://CRAN.R-project.org/package=lme4breeding), which is an open-source and powerful statistical software. Additionally, a few functions for computing relationship matrices and geospatial kernels are also included.

## Supporting information

review1

## 5 Supplementary Information

Not applicable

## 6 Acknowledgements

Special thanks to Douglas Bates, Ben Bolker and the **lme4** development team for making available the **lme4** engine for these types of extensions and answering requests and questions actively. I also want to thank Johan Aparicio-Arce and Chris Gaynor for letting me bounce some ideas with them to improve the software. I also thank Alaine Gulles for running the Asreml scripts.

## 7 Authors’ contributions

G.C.P conceived the idea, wrote the main manuscript text, prepared tables, and reviewed the manuscript.

## 8 Funding

This work was developed under the Bill and Melinda Gates Foundation ID: INV-008226 / OPP1194925.

## 9 Availability of data and materials

All programs and data sets used are available within the package at the R CRAN archive (http://cran.r-project.org/) as an open source code and can be accessed by using the help pages.

## 10 Declarations

### Ethics approval and consent to participate

Not applicable

### Consent for publication

Not applicable

### Competing interests

The author declares no competing interests.

## 11 Availability and requirements

Project name: lme4breeding Project home page: https://github.com/covaruber/lme4breeding Operating system(s): Platform independent Programming language: R Other requirements: R > 3.5 License: GPL-3 Any restrictions to use by non-academics: None

## Notes

### Competing Interest Statement

The authors have declared no competing interest.

### Summary of Updates

I have updated examples and functions. This version will be submitted to the R journal.

