## Supplementary material for "lme4breeding: a fast linear mixed model for multi-trait/multi-environment experiments": review1

**Response to reviewers**

Dear editor, I appreciate the time that you and two anonymous reviewers took to enhance the quality of this manuscript. Please find below the response to reviewers point by point.

**Reviewer #1**

Dear reviewer, I appreciate the suggestions. I have tried to incorporate your suggestions with some caveats:

1. Include lmekin and SOLAR in comparison
    a. I have tried to include the lmekin function from the coxme package in the comparison but the software doesn't allow to have both, covariances between the levels of the random effects and a covariance structure for random intercepts (e.g., environments, traits, etc.) at the same time. Only one or the other. Still, I have included lmekin in the script but warning the user that only simple models can be used. For example, a model of the type:
        i. lmekin(pheno ~ (1| id) , varlist=K, data=DT) is possible but
        ii. lmekin(pheno ~ (0+e1+e2| id) , varlist=K, data=DT) is NOT possible which is the focus of the research (multi-trait and multi-environment models with dense relationship matrices).
    b. I have also tried to include the use of SOLAR in the comparison using the R package 'solarius' but the package was archived 4 years ago. Is not compatible with new R versions and is not documented. I cloned the package from github (remotes::install_github("ugcd/solarius")) but the package doesn't run due to outdated functions that can't be debuged since they are not documented.
    c. These issues have been added to the manuscript for users to be aware that the comparison with these two software was tried.

2. For demonstration, if possible, expand the types of cases. In particular include cases where competing packages are known to perform well.
    a. I have included the script to recreate the analysis and encourage the readers to check the vignettes where multiple scenarios are extensively tested (https://CRAN.R-project.org/package=lme4breeding ). Going beyond this may expand the software note too much. I hope including all the examples in vignettes and associating them to the CRAN repository is OK with you.

**Reviewer #2**

Dear reviewer, I appreciate your suggestions. I have done my best to address the suggestions

1. As you pointed out, some sections were shortened to stay in the length expected of a software manuscript so I didn't want to expand much but I appreciate the request and opportunity to expand on clarity for the methods used in the implementation. I have

increased the length of the Implementation section to provide more clarity on the computational method and derivations together with the motivation of this additional rotation, and why this is especially useful in the multi-trait and multi-environment models.

2. Addressed in point 1. I have phrased the implementation in terms of the problem presented; multi-trait and multi-environment datasets.
3. I have added an explanation to the manuscript that since the Cholesky factor L is still dense in the lower triangular positions the system of equations is still too dense and slow to solve by lme4 compared to the classical Henderson's method. Applying the Cholesky factor to D (eigen values) instead of A makes the system fully sparse resulting in massive gains. Thanks for the request/suggestion to provide more clarity.
4. On the minor suggestions:
    a. Line 51: Should "Camel et al." be "Caamal-Pat et al."? Addressed, thanks.
    b. Line 76: What does "row-bind multiple factors" mean? Corrected, I meant row-bind multiple random effects. I have reworded.
    c. Line 78: $\sigma\sigma uu2$ is a variance component, $\boldsymbol{AA}\sigma\sigma uu2$ is the variance-covariance matrix. I reworded the paragraph to say the variance component is the variance-covariance components, and A the relationship between levels of the random effect.
    d. Line 96: After the first equality, $\sigma\sigma uu$ 2 should be of normal size (instead of a superscript); in the third equality, the second $\boldsymbol{ZZ}*$ should be $\boldsymbol{ZZ}*'$. My bad, it has been corrected for both.
    e. Line 120-122: What is the difference between $\boldsymbol{ZZ}uu$ and $\boldsymbol{ZZ}rr$? "the incidence matrix for the individuals" and "the incidence matrix for the records for the individuals" seem to be the same. Another point here is that it was stated that $\boldsymbol{HH}$ is the matrix of eigenvectors, how can it be derived as this? I would think these are two different matrices. Thanks for pointing this out, I noticed that it was not clear indeed what these two matrices mean. I have reworded and explain further how to derive them.
    f. Line 164 and lines 172-173: "$\phi\phi$ represents a one-by-one matrix of residual variance" sounds a bit strange (although mathematically it is correct). I suggest to simplify the statement as "$\phi\phi$ is the residual variance" and change "$\boldsymbol{II}2396 \otimes \phi\phi$" to "$\boldsymbol{II}2396\phi\phi$". The same for line 210, and lines 218-219. Thanks for the suggestion, this has been added.
    g. Line 195-196: I would suggest to mention the processor frequency and the number of CPUs/cores. Added.
    h. Line 210: Why was it $\boldsymbol{II}4130$? Eight traits times 413 hybrids = 3304. This is a typo and it has been corrected. Thanks for pointing that out.
    i. Line 246: … can be observed by calling the summary() function …
    j. Line 261: … in the same way as lme4 since …
    k. Table 1: Please be consistent with respect to the form of the decimals. For example, in the sixth column, there were "0.04" and "-.07". It has been corrected, all were followed by a zero so I ignored it but I agree with the suggestion.
    l. Table 2: What is the different between "Sommer-mmec" and "sommer" (in Table 1)? I have added an explanation below Table 1.

m. Table 3: Why only two software were compared for this data set? Since unstructured models have a hard time to converge when the number of traits increases, the use of the other software was prohibitive due to computation time and singularities during the REML process. I have added an explanation in the implementation section to make readers aware.
